# Semi-synthetic Cinnamodial Analogues: Structural Insights into the Insecticidal and Antifeedant Activities of Drimane Sesquiterpenes Against the Mosquito *Aedes aegypti*

**DOI:** 10.1101/536961

**Authors:** Preston K. Manwill, Megha Kalsi, Sijin Wu, Xiaolin Cheng, Peter M. Piermarini, Harinantenaina L. Rakotondraibe

## Abstract

The *Aedes aegypti* mosquito serves as a major vector for viral diseases, such as dengue, chikungunya, and Zika, which are spreading across the globe and threatening public health. In addition to increased vector transmission, the prevalence of insecticide-resistant mosquitoes is also on the rise, thus solidifying the need for new, safe and effective insecticides to control mosquito populations. We recently discovered that cinnamodial, a unique drimane sesquiterpene dialdehyde of the Malagasy medicinal plant *Cinnamosma fragrans,* exhibited significant larval and adult toxicity to *Ae. aegypti* and was more efficacious than DEET – the gold standard for insect repellents – at repelling adult female *Ae. aegypti* from blood feeding. In this study several semisynthetic analogues of cinnamodial were prepared to probe the structure-activity relationship (SAR) for larvicidal, adulticidal and antifeedant activity against *Ae. aegypti*. Initial efforts were focused on modification of the dialdehyde functionality to produce more stable active analogues and to understand the importance of the 1,4-dialdehyde and the α,ß-unsaturated carbonyl in the observed bioactivity of cinnamodial against mosquitoes. This study represents the first investigation into the SAR of cinnamodial as an insecticide and repellent against the medically important *Ae. aegypti* mosquito.

## Introduction

Mosquitoes are vectors of numerous human pathogens, such as the malaria parasite, dengue virus, chikungunya virus, and Zika virus, which affect over 300 million people annually^1–3^. While the majority of the burden has been shouldered by Africa and South-East Asia the global disease distribution is widening. The worldwide incidence of dengue has risen 30-fold in the past 30 years, and more countries are reporting their first outbreak of the disease^3^. Chikungunya and Zika viruses, both historically limited to parts of Africa and Asia, have recently emerged into global threats with increased transmission in the Americas^4,5^. The arboviruses that cause dengue, Zika, chikungunya and yellow fevers are all transmitted to humans by the mosquito *Aedes aegypti*. According to the World Health Organization more than half of the world’s population lives in areas where this mosquito species is present, including several southern regions in the United States^2^. While significant progress has been made in developing therapeutics and vaccines for mosquito-borne pathogens, more effective and low-cost means to treat and prevent these diseases are still underdeveloped or unavailable^6,7^. Vector control strategies remain the primary method to control and prevent the spread of mosquito-borne diseases^8^; chiefly, control of mosquitoes with insecticides is often the only method proven to reduce vector populations during an emerging epidemic^9^.

The major classes of insecticides used in vector control strategies include the pyrethroids, carbamates, organophosphates, and neonicotinoids, which all target the nervous system of insects^10–13^. While their activity has made them very effective at reducing mosquito populations, they are non-selective, killing beneficial insects and in some cases small vertebrate animals, which has caused the removal of some agents such as DDT and other organochlorine compounds from the vector control arsenal^14^. Excessive use of the remaining groups of insecticides, however, has led to the selection of insecticide-resistant mosquito populations^15–17^. Moreover, no new public health insecticides have been developed in the past 40 years^18^. Thus, it is imperative that we replenish our chemical toolbox by identifying new agents that exhibit novel mechanisms of action with high selectivity to mosquitoes.

Plants have been an indispensable source of novel compounds possessing pharmacological activities relevant to public health^10^. Pyrethroids, for instance, the most widely used insecticides in the United States and the only class approved for insecticide treated nets^19^, are derived from natural pyrethrins isolated from the flowers of *Chrysanthemum* (Asteraceae)^20^. Recently, we have identified that an extract of *Cinnamosma fragrans* Baill. (Canellaceae), a plant used in Malagasy traditional medicine, is antifeedant, repellent, and toxic to *Ae. aegypti* mosquitoes. In our efforts to isolate and characterize the bioactive compounds from *C. fragrans*, we identified cinnamodial (CDIAL, **1**), a drimane sesquiterpene with promising toxicity to larval and adult female *Ae. aegypti* mosquitoes^21^. In addition to exhibiting a similar toxic profile against pyrethroid-susceptible and -resistant strains of *Ae. aegypti*, CDIAL was more efficacious than DEET [N,N-Diethyl-meta-toluamide] – the gold standard for insect repellents – at repelling mosquitoes from feeding on blood^21^. Moreover, we demonstrated that the mechanism of the antifeedant activity of CDIAL was through the activation of transient receptor potential A1 (TRPA1) channels^21^.

The goal of the present study was to investigate the structural basis of the insecticidal and antifeedant activities of **1** against *Ae. aegypti* by generating a series of semi-synthetic CDIAL derivatives. Our efforts led to the discovery of **10** ((-)-6*ß*-Acetoxy-9*α*-hydroxydrim-7-ene-12-methyl-12-one-11-al) as a CDIAL derivative with superior and similar insecticidal activity against larvae and adult females, respectively, but weaker antifeedant activity against adult females. Herein, we describe the re-isolation of **1**, analog generation, and biological evaluation of synthetic derivatives. Additionally, in order to understand the bioactivity of the most active compounds, the observed structure-activity relationship (SAR) and potential reaction of the active compounds with primary amines of mosquito TRPA1 are discussed.

## Results and Discussion

### Isolation and Identification of Bioactive Drimane Sesquiterpenes

The powdered stem bark of *C. fragrans* (Canellaceae) was extracted with dichloromethane, concentrated, and subjected to silica gel column chromatography and recrystallization to afford cinnamodial (**1**).

### Derivatization of Cinnamodial

Cinnamodial (**1**) is one of the approximately more than 80 naturally occurring terpenoids containing an α,β-unsaturated 1,4-dialdehyde functionality^22^, specifically **1** belongs to the drimane sesquiterpene class of compounds which includes the structurally similar compounds: warburganal (**2**) and polygodial (**3**) (**Fig. 1**). Unsaturated dialdehyde-containing compounds exhibit diverse bioactivities^23,24^, including antimicrobial^25^, antifungal^25^, molluscicidal^26^, and cytoxicity^27^. Additionally compounds **1**, **2** and **3** are pungent to humans^28–30^, possess antifeedant and insecticidal activity^31,32^, and agonize Transient Receptor Potential A1 (TRPA1) channels^21,22,30,33,34^. Most biological activities, including the antifeedant activity, of these drimane sesquiterpenes have been attributed to the α,β-unsaturated 1,4-dialdehyde functionality of the molecules, forming adducts with free sulfhydryl groups^35,36^ or primary amines such as the ε-amino group of lysine^37,38^. Our previous results showed that **1** could effectively kill mosquito larvae in an aqueous environment, penetrate the cuticle of adult female mosquitoes, reduce the feeding of mosquitoes when added to a sucrose solution, and reduce the propensity of mosquitoes to blood feed when dried onto the surface of a membrane feeder. Despite these promising results, nonspecific reactions of dialdehydes with endogenous free amines or water may reduce the bioavailability of these compounds. In this work, initial semi-synthetic modification was focused on producing more stable derivatives by replacing the two aldehyde groups while maintaining the α,β-unsaturated system.

**Figure 1.**
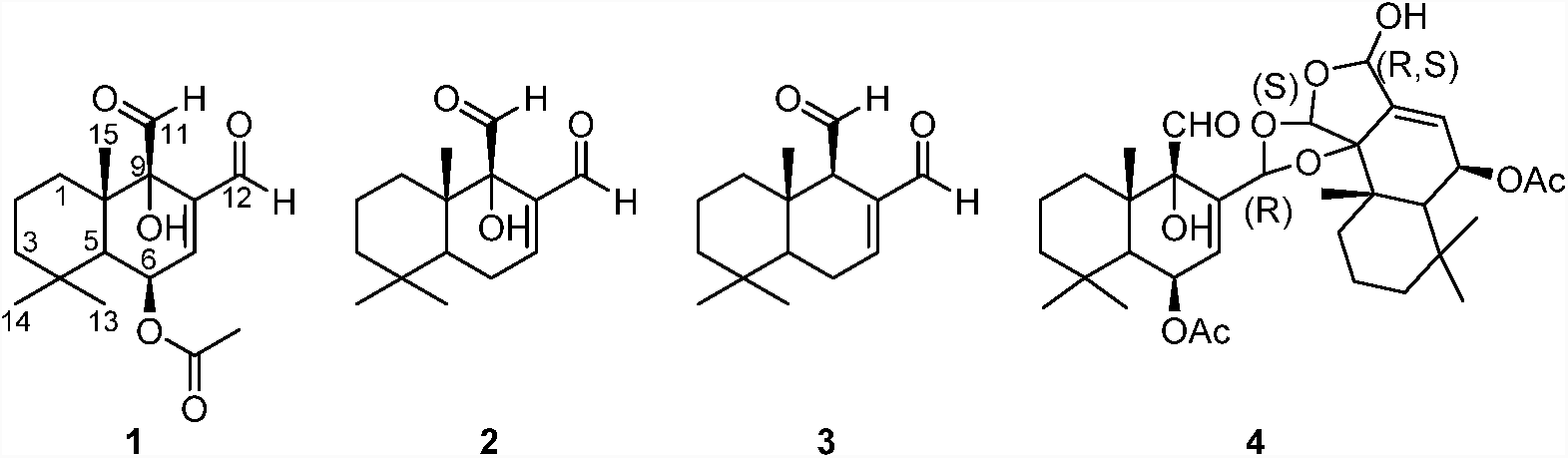
Chemical structures of natural drimane-type compounds including those isolated from *C. fragrans* (**1**, **3**, and **4**): Cinnamodial (**1**), Warburganal (**2**), Polygodial (**3**), Capsicodendrin (**4**).

Cinnamodial **1** was used as starting material for derivatization and analog generation (**Scheme 1**). Pinnick oxidation was employed to generate the cinnamodiacid (**5**), wherein the C-11 and C-12 aldehydes were successfully oxidized to the corresponding carboxylic acids. The methyl ester derivative (**6**) was obtained by reacting **5** with (Trimethylsilyl)diazomethane. Reaction of **1** with hydrazine afforded a 2,3-dihydro-3-pyridazinol (**7**), which maintained the unsaturated system.

**Scheme 1.**
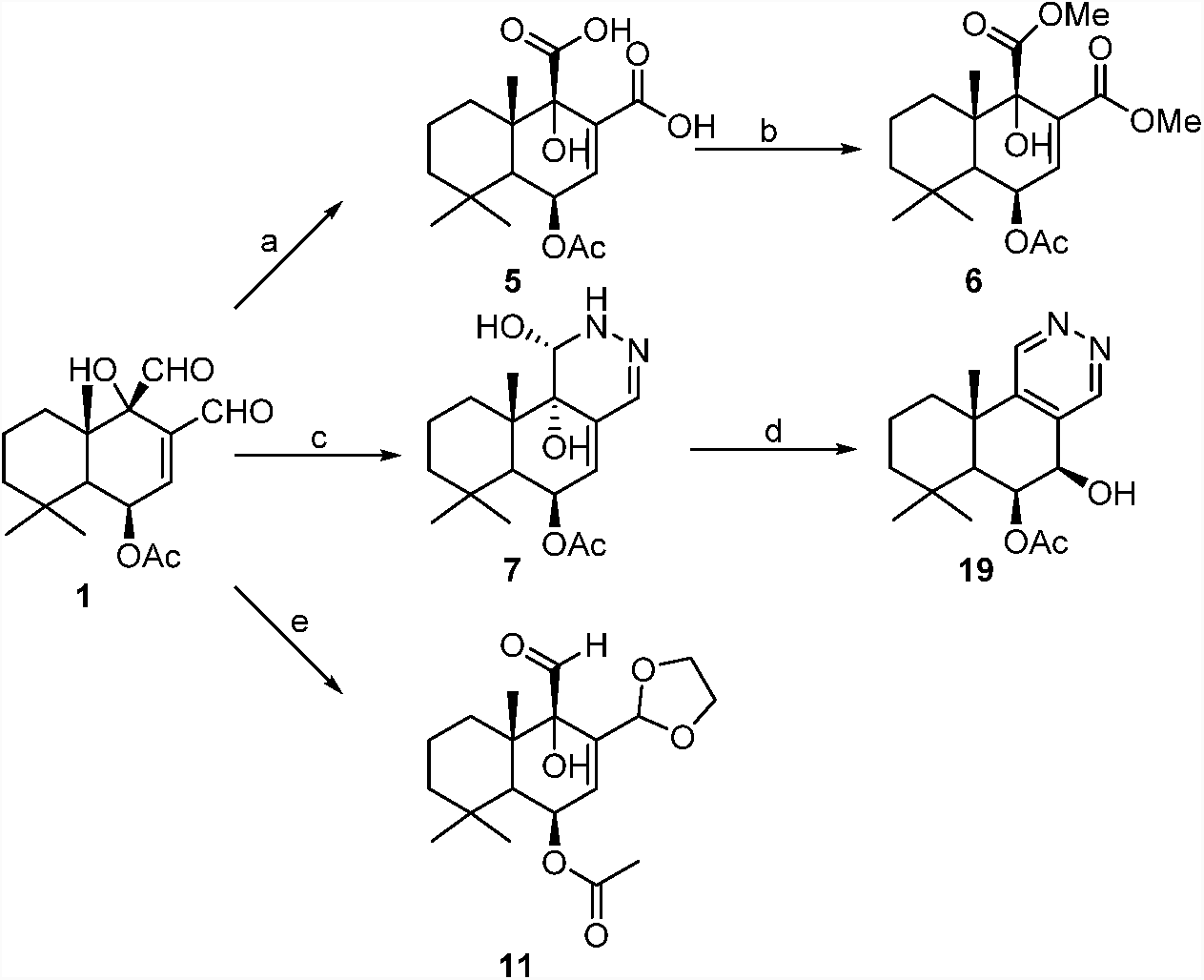
Derivatization of **1** into structural analogues. Reagents and conditions: (a) NaClO_2_, NaH_2_PO_4_·H_2_O, H_2_O_2_, MeCN, H_2_O, r.t., 24 h; (b) TMS-CHN_2_, MeOH:Et_2_O, 0°C, 30 min;(c) H_2_N-NH_2_, CH_2_Cl_2_:EtOH, 2 h; (d) MeOH, r.t.; (e) HOCH_2_CH_2_OH, *p*-TsOH, benzene, reflux, 24 h.

It was envisioned that the more reactive C-12 aldehyde of **1** could be selectively transformed into a methyl ketone by treatment with a methyl Grignard or methyl lithium reagent followed by oxidation of the resulting secondary alcohol. Treatment of **1** with one equivalent of methylmagnesium bromide or methyl lithium instead produced two diastereomeric lactols (**8**) and (**9**), instead of a secondary alcohol. As determined by NMR studies (see experimental), the lactols **8** and **9** differ at the C-12 position depending on whether the C-12 aldehyde of **1** was attacked from the α or ß face, respectively (**Scheme 2**). Lactol **9** was then converted into the desired compound (**10**), with a C-11 aldehyde and C-12 ketone, upon incubation in deuterated chloroform. However, the isomer **8** remained stable in deuterated chloroform and even resisted base-induced lactol-opening / tautomerization with 1,8-Diazabicyclo(5.4.0)undec-7-ene (DBU), see supporting information.

**Scheme 2.**
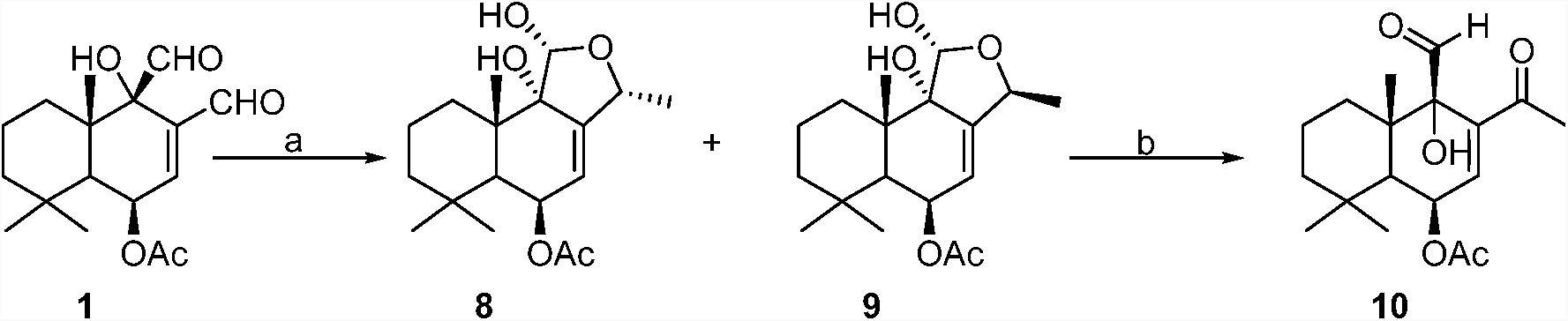
Synthesis of lactols **8** and **9**, and formation of **10** from lactol **9**. Reagents and conditions: (a) 1 eq. MeMgBr, THF, −78°C, 2 h; (b) CDCl_3_, 22 d.

To understand the role of the unsaturated system and confirm the necessity of the C-12 unsaturated aldehyde for its observed biological activity^21^, **1** was exposed to one equivalent of ethylene glycol and a catalytic amount of *p*-toluenesulfonic acid in benzene to afford the monoacetal **11** (**Scheme 1**).

Plants and marine organisms have been shown to produce a number of cytotoxic quinones and hydroquinones, including avarone, bolinaquinone, juglone, and lapachol^39,40^. Lapachol, isolated from *Tabebuia avellanedae*, and structurally related derivatives have shown modest larvicidal activity against mosquitoes^41,42^. Further modifications of **1** were implemented to prepare a derivative potentially exhibiting mosquito toxicity without antifeedant activity. We had previously shown that the carbon at C-9 bearing a hydroxyl in capsicodendrin (**4**), could be transformed into a ketone by treatment with pyridinium chlorochromate^43^. Therefore, in order to produce a dihydroquinone, **1** was reacted with 5 equivalents of the MeMgBr to alkylate the C-12 aldehyde and to cleave off the C-6 acetyl group providing an isomeric mixture of deacetylated lactols 12α-methyl-pereniporin A and 12ß-methyl-pereniporin A (**12**) (**Scheme 3**). Pyridinium chlorochromate in dichloromethane was then used to oxidize **12** into a mixture of C-12 isomers of (1′S/R)-1′-((8aS)-5,5,8a-trimethyl-1,4-dioxo-1,4,4a,5,6,7,8,8a-octahydronaphthalene-2-yl)ethyl formate (**13**).

**Scheme 3.**
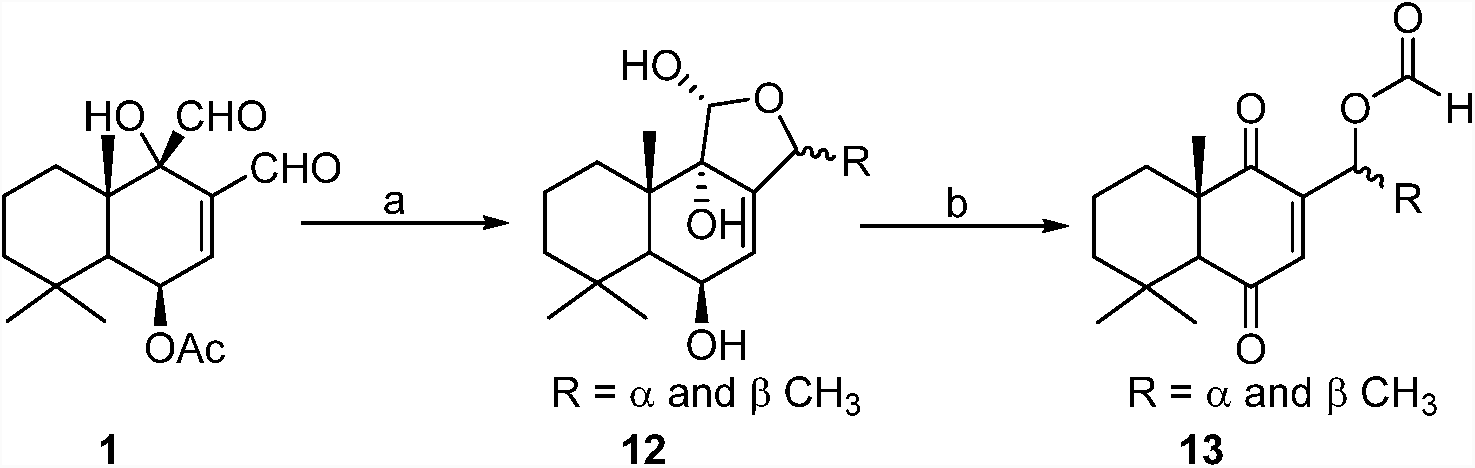
Synthesis of 1,4-dione **13**. Reagents and conditions: (a) 5 eq. MeMgBr, THF, −78°C, 2 h; (b) 3.3 eq. PCC, DCM, 0 °C → r.t., 2h.

### Model Study on the Nucleophilic Addition of Thiol or Amino Residues to Cinnamodial

Studies on natural sesquiterpenes containing the α,β-unsaturated 1,4-dialdehyde moiety, namely, polygodial (**3**), miogadial (**14**), and isovelleral (**15**) (**Fig. 2**), have suggested that these molecules activate TRPA1 through a mechanism different from that of reactive α,β-unsaturated aldehydes and isothiocyanates^44–48^. Specifically, α,β-unsaturated aldehydes such as the endogenous ligand (4-hydroxynonenal), the main odiferous compound in cinnamon (cinnamaldehyde), and the irritant compound acrolein, activate the TRPA1 channel by covalent modification of N-terminal cysteine residues^49–52^. Sesquiterpenes with a α,β-unsaturated 1,4-dialdehyde moiety, on the other hand, have been shown to undergo Paal-Knorr condensation reactions with lysine residues^30,34,53^. Since there have been no studies showing the reaction of CDIAL with thiol or amine residues, the reaction of L-cysteine methyl ester with CDIAL was used as a model system to study the reaction of **1** with a biological substrate which may in principle react by either thiol or amine addition. CDIAL was treated with L-cysteine methyl ester under basic conditions (in pyridine) to afford the pyrrolidine (**16**). This adduct likely formed from the initial attack by the primary amine at the more reactive C-12 aldehyde to give the azomethine (**A**) intermediate, which was then attacked by the thiol to form pyrrolidine (**16**) (**Scheme 5**). The reaction product and proposed reaction mechanism suggest that the α,β-unsaturated 1,4-dialdehyde (**1**) may also activate TRPA1 by forming reactive pyrrole-type conjugates with the amino groups present in the protein, as demonstrated for warburganal **2**, polygodial **3**, and 1ß-acetoxy-9-deoxy-isomuzigadial (**17**)^30^. Additionally, the attack of the azomethine (**A**) by the thiol of L-cysteine methyl ester suggests that the reactive imine can be attacked by nearby nucleophilic groups, such as the thiol of cysteine and hydroxyl groups of serine or threonine.

**Figure 2.**
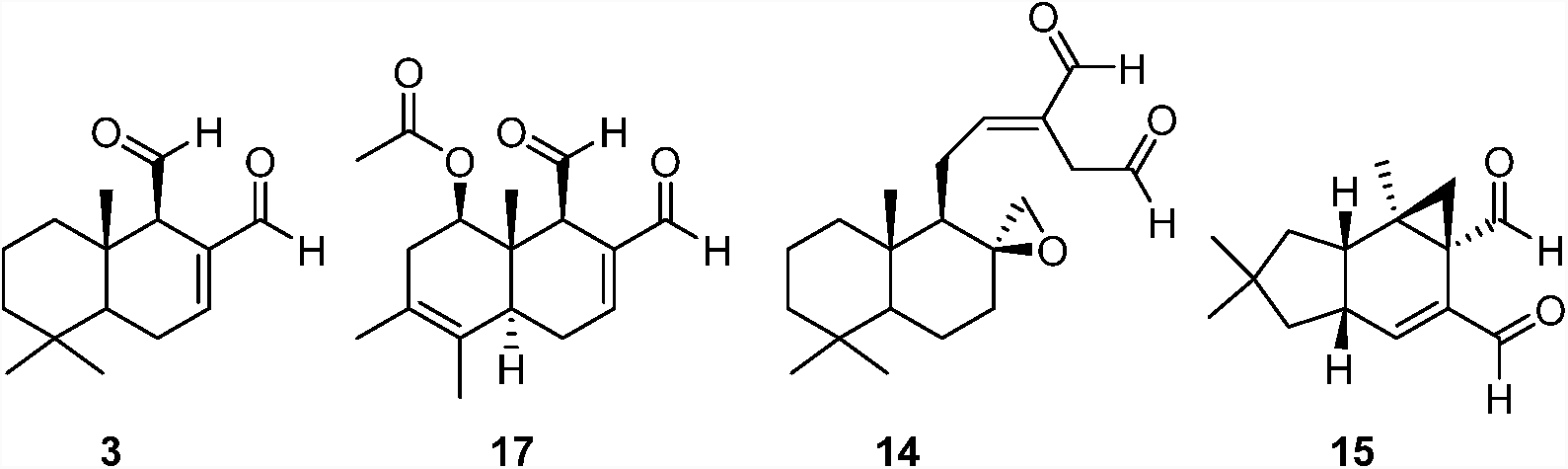
Chemical structures of α,ß-unsaturated 1,4-dialdehydes: polygodial (**3**), 1ß-acetoxy-9-deoxy-isomuzigadial (**17**), miogadial (**14**), (+)-isovelleral (**15**).

**Scheme 4.**
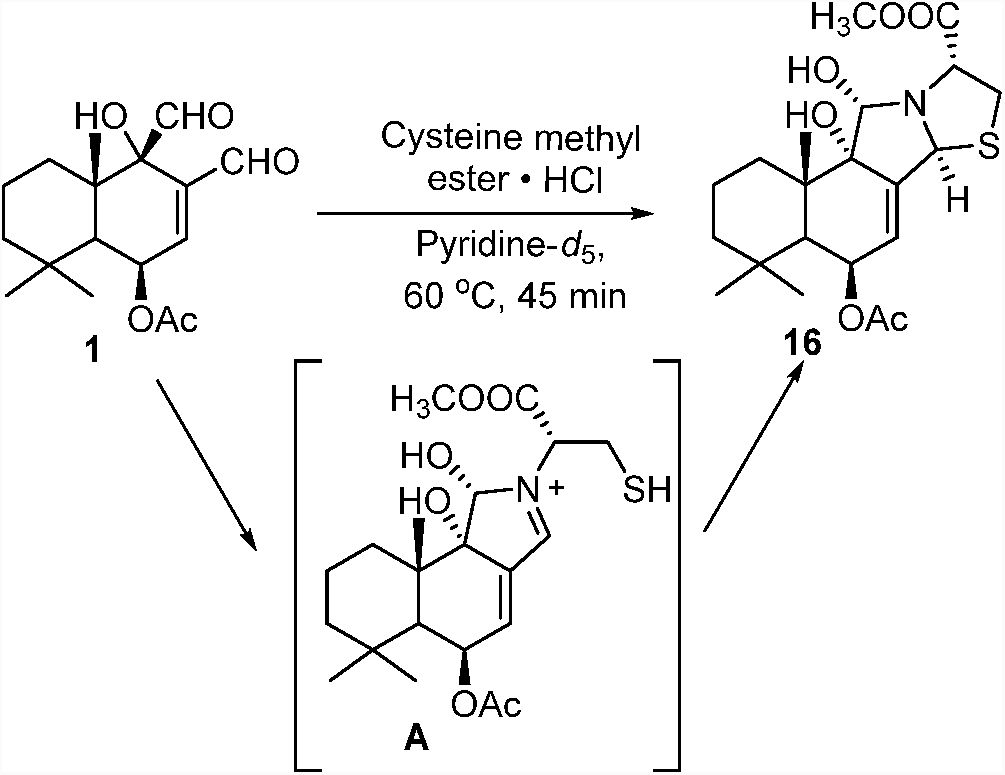
Reaction mechanism of the reaction of **1** with L-cysteine methyl ester via a cationic azomethine (**A**) to form the pyrrolidine (**16**).

### Toxicity to larval and adult female mosquitoes

All prepared CDIAL derivatives were screened for 24 h larvicidal activity against 1^st^ instar *Ae. aegypti* (Liverpool, LVP, strain) using a concentration of 100 µM in the rearing water^21^. At this concentration, CDIAL killed *ca.* 70% of the larvae within 24 h (**Fig. 3A**). Nearly all of the CDIAL derivatives showed significantly lower efficacy than CDIAL. However, compounds **10** (86.5%) and **13** (100%) exhibited statistically greater efficacy than CDIAL (**Fig. 3A**).

**Figure 3.**
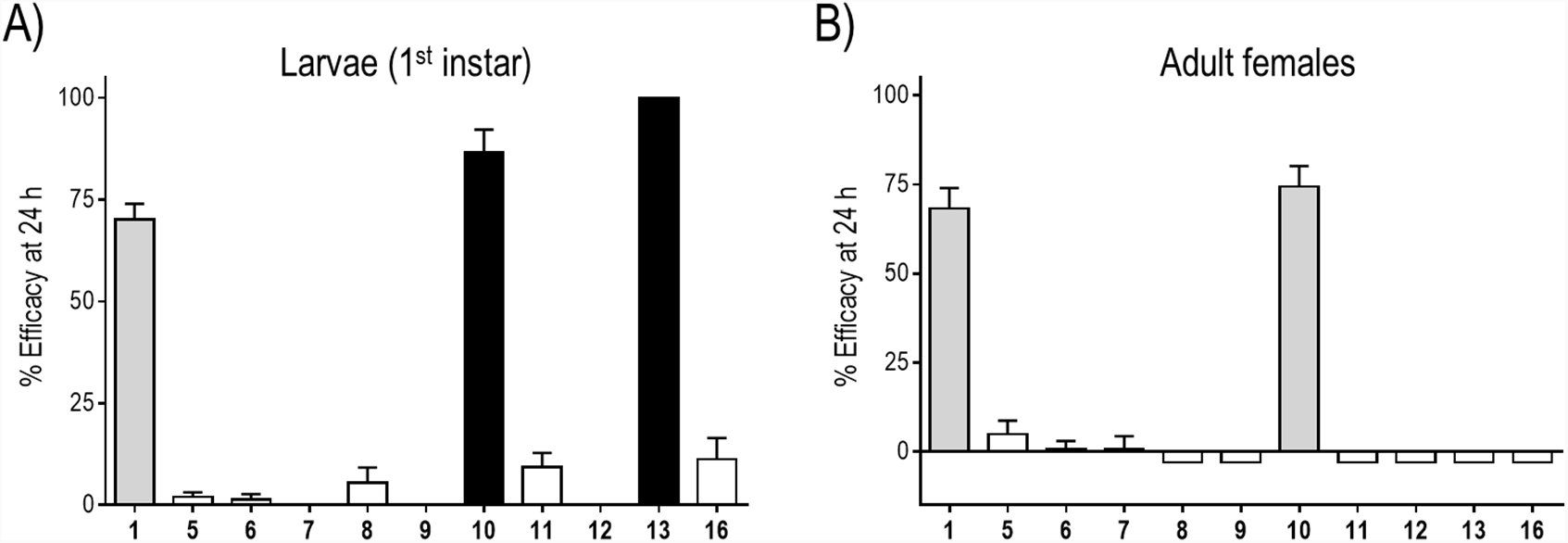
Toxic efficacy of cinnamodial and derivatives against larval (A) and adult female (B) *Ae. aegypti*. The efficacy values were calculated using Abbott’s correction to account for control (100% acetone) mortality^56^. In A), compounds were added to the rearing water of 1^st^ instar larvae (100 µM) and efficacy was defined as the percentage of larvae that died within 24 h. In B), compounds were applied to the thoracic cuticle of adult females (1.5 nmol/mosquito) and efficacy was defined as the percentage of adults that were incapacitated (dead or flightless) within 24 h. Values are means ± SEM based on at least 6 replicates of 5 larvae each or 3 replicates of 10 adult females each. Shading indicates statistical categorization of derivative’s efficacy relative to CDIAL as determined by a one-way ANOVA with a Bonferroni post-test: gray = similar (P > 0.05); filled = superior (P < 0.05); open = inferior (P < 0.05).

The derivatives were also screened for 24 h toxicity against adult female *Ae. aegypti* using a dose of 1.5 nmol applied to the thoracic cuticle of each mosquito. At this dose, CDIAL incapacitated 68% of the mosquitoes within 24 h. All of the CDIAL derivatives were significantly less efficacious than CDIAL except for **10**, which was similar to CDIAL in efficacy (74%).

Although the acid **5**, the ester **6**, and the 2,3-dihydro-3-pyridazinol **7** derivatives maintained the α,β-unsaturated functionality of **1**, they were inactive in the toxicity assays against larvae and adult females. The lack of toxicity of **7** can be due to its conversion to the 7-hydroxy-pyridazine (**18**), which has been observed in deuterated methanol at room temperature, and also may occur in the carrier solvent used for the bioassays or upon exposure to the rearing water. Since lactol derivatives (**8**, **9**, and **12**) were not toxic to larvae and adult females, we conclude that the drimane skeleton with C-11 hemiacetal polar head groups is insufficient to elicit mosquito toxicity. These results are consistent with published findings that the biological activity of drimane-type compounds are reduced or lost when the aldehyde functionalities are modified^31,54,55^. Acetal protection of the C-12 aldehyde as in compound (**11**) also abolished the toxicity to larvae and adult females, thus illustrating the crucial role of the conjugated C-12 aldehyde in eliciting a biological response. On the other hand, compound **10**, a C-12 methyl ketone of **1**, exhibited superior and similar efficacy to larvae and adult females, respectively, relative to **1**. Thus, the more stable methyl ketone can effectively replace the aldehyde moiety. Remarkably, the dihydroquinone (**13**) exhibited superior larvicidal activity to **1**, but was nominally toxic to adult females. We suspect that the chemical modifications that result in the dihydroquinone preserve the toxicity of the molecule, but may reduce its ability to penetrate the cuticle of adult females. In summary, the relative activities of the analogs observed against larval and adult mosquitoes (**Fig. 3**) indicate the important role of the 1,4-dialdehyde functional group in insecticidal activity.

### Antifeedant activity in adult female mosquitoes

A capillary feeding (CAFE) choice bioassay was used to screen the antifeedant activity of the CDIAL derivatives against adult female *Ae. aegypti* (LVP strain)^21,57,58^. Briefly, mosquitoes were presented with two capillaries of 10% sucrose as a food source for 18-20 h; the ‘control’ capillary was treated with 1% DMSO (the carrier solvent for **1** and its derivatives in this assay) while the ‘treatment’ capillary was treated with **1** or a derivative thereof at a concentration of 1 mM. All of the derivatives elicited inferior antifeedant activity compared to CDIAL (**Fig. 4**).

**Figure 4.**
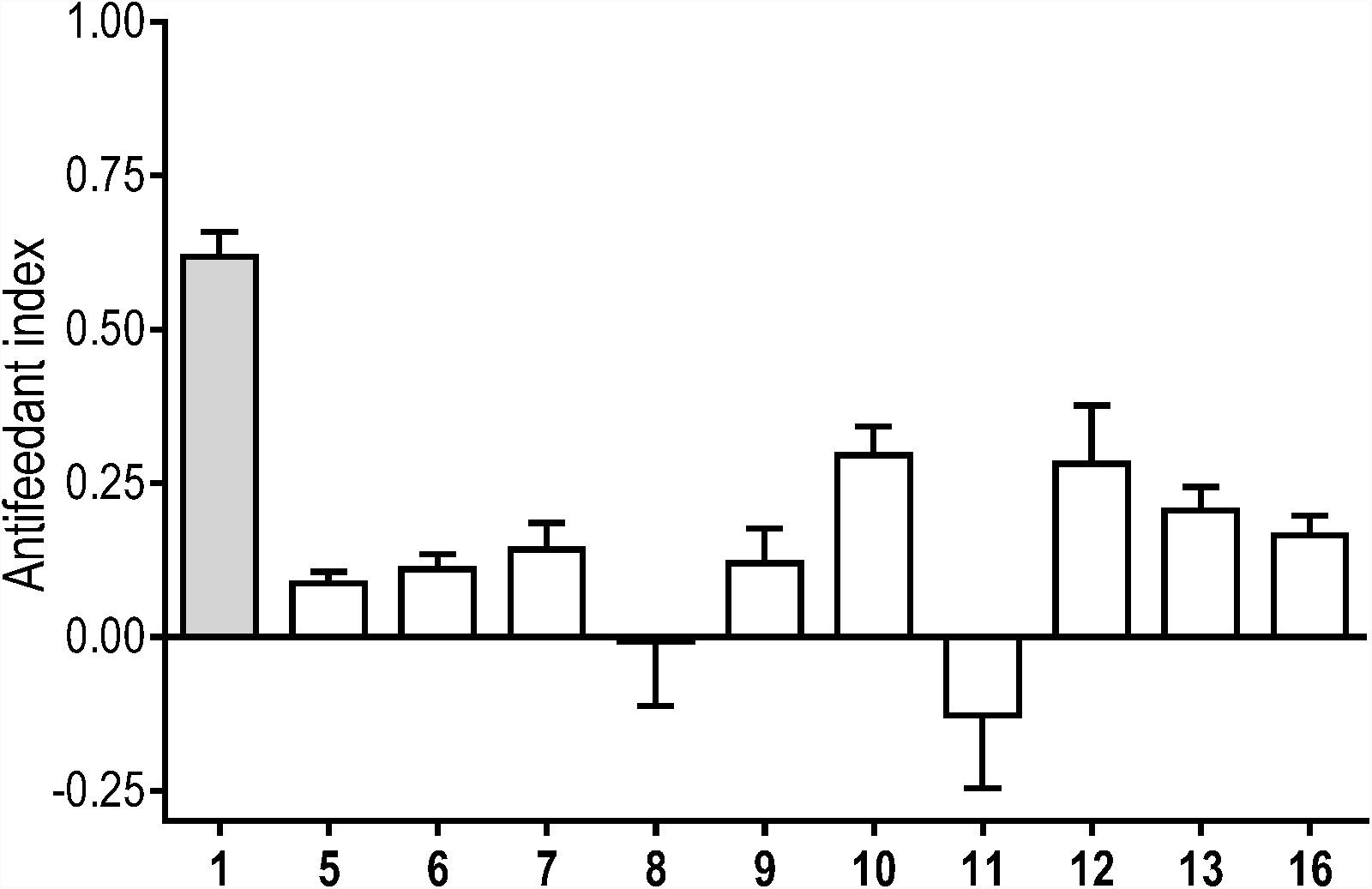
Antifeedant activity of CDIAL and semi-synthetic derivatives as determined via choice CAFE assays in adult female *Ae. aegypti* (LVP strain). Mosquitoes were allowed to feed equally on two capillaries of 10% sucrose with 0.01% trypan blue; the control capillary included 1% DMSO, and the treatment capillary included 1% DMSO and 1mM test compound. The difference in volume consumed between the capillaries was used to calculate the antifeedant activity^**^34^**^. Values are means ± SEM based on at least 8 replicates of 5 adult females each. Shading indicates statistical categorization of derivative’s efficacy relative to CDIAL as in **Figure 3**.

We have previously shown that the antifeedant activity, but not insecticidal activity, of CDIAL was associated with its modulation of TRPA1 channels^21^. Thus, we suspect that all of the derivatives were inferior agonists of TRPA1 compared to CDIAL. The reduced TRPA1 agonistic activity may be attributable to the reactivity of the unsaturated system of the molecules. Specifically, the unsaturated aldehyde in **1** favors 1,2-addition, while **5**, **6**, and **7** likely favor 1,4-addition. Thus, **1** would be more amenable to direct attack of an amino group on TRPA1 than the other conjugated derivatives. The mosquito antifeedant activities of **5**, **6**, and **7** showed a similar trend to previously reported antifeedant activity of polygodial analogues^23,24^. As mentioned above, the possible conversion of **7** to **18** during the assay may also lead to the reduced activity. The absence of activity in **11** illustrates the necessity of the C-12 conjugated carbonyl group.

Although the derivatives were inferior to CDIAL in the context of antifeedant activity, bioactivity was not completely lost and they still provide some insights into the SAR. For example, among the derivatives, **10** and **12**, were relatively effective antifeedants. The moderate activity of compound **10** may indicate that the C-12 aldehyde can be converted into a methyl ketone without complete loss of antifeedant efficacy. For the lactol **12** it is possible that a ring opening tautomerization affords the C-11 aldehyde and a C-12 conjugated carbonyl.

Unsaturated aldehyde-containing agonists, such as acrolein, are known to react with nucleophilic residues of TRPA1 channels, including cysteine, histidine, and lysine. Several cysteine residues in TRPA1 channels have been identified as potential sites that can form covalent adducts with these agonists^**49,50**^. However, further mutagenesis experiments also showed that unsaturated dialdehyde-containing sesquiterpenes might not bind to the same sites as those targeted by the small agonists^**47**^. To shed light on how CDIAL interacts with the mosquito TRPA1 channel and thereby modulating its antifeedant effects, we have developed a structural model of mosquito TRPA1 based on the single-particle cryo-electron microscopy (EM) structure of human TRPA1 (hTRPA1)^**59**^, and then computationally docked CDIAL to the AgTRPA1 structural model. We used the TRPA1 channel of *Anopheles gambiae* (AgTRPA1) as a representative mosquito TRPA1, because it has been previously cloned and shown to be directly activated by CDIAL^**21,60**^. Moreover, the amino acid identity between AgTRPA1 and the predicted *Ae. aegypti* TRPA1 (AAEL009419) is very high (>83%). In particular, the putative CDIAL binding region we have identified (see below) is over 90% identical between the two species.

Our results suggest that CDIAL binds preferentially to a ‘pocket’ near Cys684 (**Fig. 5**), a residue in TRPA1 channels that have been implicated to interact with electrophilic agonists^**50**^. Notably, six lysine residues are located within a radius of 10 Å from Cys684, including Lys656, Lys678, Lys681, Lys728, Lys738 and Lys744. Many of these lysine residues are surrounded by negatively charged residues (Glu or Asp), thereby making them more nucleophilic and likely to interact with an electrophile like CDIAL. The dominant predicted CDIAL binding site is located between Cys684 and Lys728 (**Fig. 5**). Analogous to Lys661 in human TRPA1, Lys728 is the closest lysine to Cys684. In these docked poses (**Fig. 5**), the C-12 aldehyde of CDIAL is found close (4.2-5.0 Å) to the amino group of Lys728 that can initiate a nucleophilic attack on C-12 to form an initial CDIAL-AgTRPA1 conjugate, highlighting the importance of the reactive C-12 aldehyde group. This is consistent with the observation that acetal protection of the C-12 aldehyde in compound 11 abolished its antifeedant activity (**Fig. 4**). Further, given the proximity (∼5.0 Å) of the thiol group of Cys684, the polarized C-12 or C-7 of CDIAL may subsequently be attacked by this thiol. Overall, the docking results show a few possible ways in which CDIAL can interact with AgTRPA1, suggesting that the reactions of the α,β-unsaturated 1,4-dialdehyde with nearby amine and thiol side chains (consistent with the formation of pyrrolidine when CDIAL was treated with L-cysteine methyl ester in pyridine proposed in **Scheme 5**) may be responsible for activating the AgTRPA1 channel. Overall, these preliminary computational results provide a molecular rational for the importance of the dialdehyde moiety and the α,ß-unsaturated carbonyl for CDIAL’s antifeedant activity. Nonetheless, due to the high flexibility of the loops shaping the CDIAL binding pocket, our computational studies do not exclude the possibility that other Lys residues such as Lys681 and Lys744 may also be involved in reacting with **1** after some local conformational rearrangements.

**Figure 5.**
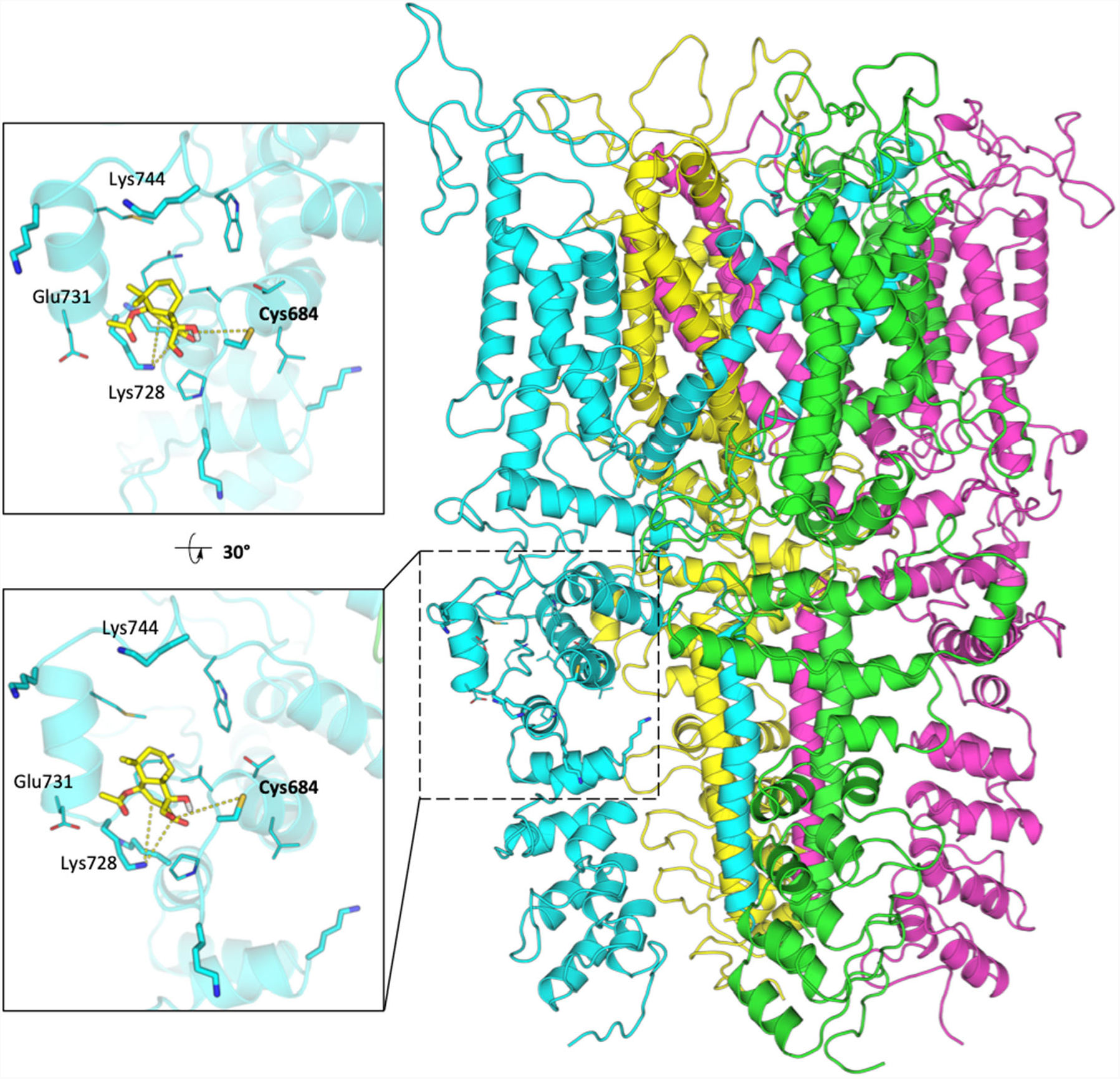
The computationally docked structure of CDIAL (**1**) in AgTRPA1. The tetrameric AgTRPA1 structure is shown on the right in a cartoon representation. Two zoomed views of the CDIAL binding site are shown on the left. The ligand is shown in a yellow licorice representation and the protein in a cyan cartoon representation. Nearby Cys and Lys residues are labeled.

## Conclusion

The current study has analyzed ten semisynthetic analogs (**5**-**13**, and **16**) of the α,ß-unsaturated 1,4-dialdehyde cinnamodial (**1**) to identify the structural contributions of **1** toward its larvicidal, adulticidal and antifeedant activity against *Ae. aegypti*. Two analogs, **10** and **13** exhibited more efficacious toxicity against mosquito larvae than **1**. Moreover, **10** was of similar toxic efficacy against adult females as **1**. These results indicate that the reactive C-12 aldehyde can be substituted with relatively stable moieties, such as an unsaturated methyl ketone (**10**) or hydroquinone (**13**), without sacrificing insecticidal activity. All of the other CDIAL analogs showed that the bicyclic drimane skeleton alone was not sufficient to induce larval or adult toxicity.

Notably, all of the analogs possessed weaker antifeedant activity than **1** regardless of their insecticidal activity. These results support the notion that CDIAL’s insecticidal and antifeedant mechanisms of action are independent^21^. While our previous results associated CDIAL’s antifeedant activity with its modulation of TRPA1^21^, this work suggests that CDIAL may interact with TRPA1 by an initial nucleophilic attack at C-12 by a lysine residue to form a CDIAL-TRPA1 conjugate. The activated conjugate may then undergo further attack at the C-12 or C-7 of CDIAL by a neighboring thiol or nucleophilic residue^38^.

Despite the thorough elucidation of the antifeedant structure-activity relationship against lepidopterans^26,32,37,61,62^ and the toxicant activity against insect pests^55,61,63–65^ by several groups, this is the first investigation into the observed structure-activity relationship (SAR) of **1** against mosquitoes (**Fig. 5**). Overall, we have identified several CDIAL derivatives that confirm the importance of the dialdehyde moiety for mosquitocidal activity and the α,ß-unsaturated carbonyl for antifeedant activity. The improved larvicidal activity observed in **10** and **13** provide insights that will aid the future development of more stable and effective insecticidal CDIAL derivatives.

### Experimental Section

#### General Experimental Procedures

Optical rotations ([α]_D_) were measured with an LED light source monitoring at 589 nm in acetonitrile at 20 °C. The instrument used was an Anton Paar MCP 150 polarimeter (Anton Paar OptoTec GmbH, Seelze-Letter, Germany). Ultraviolet (UV) absorption spectra were measured in a 1 cm quartz tank using a Hitachi U-2910 UV/vis double-beam spectrophotometer (Hitachi High-Technologies America, Schaumburg, IL, USA). Infrared (IR) spectra were obtained on a Nicolet 6700 FT-IR spectrometer (Thermo Scientific, Waltham, MA, USA). The 1D (^1^H, ^13^C, selective COSY and NOESY) and 2D (COSY, HSQC, HMBC, and NOESY) NMR spectra were recorded at 300 K (26.85 °C) in CDCl_3_ for all compounds except for **5** and **19** which were measured in methanol-*d*_4_ and **7** which was measured in acetonitrile-*d*_3_ on a Bruker Avance III HD 400 MHz instrument (Bruker, Billerica, MA, USA) using standard Bruker pulse sequences. ^1^H chemical shifts are reported in parts per million (ppm) and are referenced to the residual CDCl_3_ signal (δ 7.26 ppm), methanol-*d*_4_ signal (δ 3.34 ppm), pyridine-*d*_5_ signal (δ 7.22 ppm), or acetonitrile-*d*_3_ signal (δ 1.96 ppm). ^13^C chemical shifts are reported in ppm and are referenced to the residual CDCl_3_ signal (δ 77.36 ppm), methanol-*d*_4_ signal (δ 49.86 ppm), pyridine-*d*_5_ signal (δ 123.87 ppm) or acetonitrile-*d*_3_ signal (δ 118.77 ppm). Deuterated NMR solvents were purchased from Cambridge Isotope Laboratories (Tewksbury, MA, USA). High-resolution mass spectra (HRESIMS) were acquired on a hybrid spectrometer utilizing a linear ion trap and Orbitrap (LTQ Orbitrap, ThermoFisher Scientific Inc., Bremen, Germany) equipped with an ESI source in the positive-ion mode, with sodium iodide (NaI) being used for mass calibration. The spectrometer was equipped with an Agilent 1100 HPLC system (Agilent Technologies, California, USA) including a binary pump, UV detector, and autosampler. Data acquisition and analysis were accomplished with Xcalibur software version 2.0 (Thermo Fisher Scientific Inc., Bremen, Germany). The samples were prepared at a concentration of ∼10 μg/mL in ACN, and the injection volume was set at 30 μL. Flash and open-column chromatography were performed with SilicaFlash^®^ P60 (230-400 mesh; SiliCycle Inc., Quebec City, Canada) using solvent systems as described. Ratios of solvent systems used for chromatography are expressed in v/v as specified. Analytical thin-layer chromatography was performed on aluminum-backed precoated silica gel plates (0.24 mm; Dynamax Adsorbant, Inc., Darmstadt, Germany). Spots were visualized under UV light (254 & 320) and by spraying with modified Godin’s reagent (vanillin/EtOH-Perchloric acid, 1:1, v/v) and H_2_SO_4_-water (15%) followed by heating. Commercially available chemicals were used as purchased. Ice/water was used as the temperature bath to achieve 0 °C and dry ice/acetone was used to achieve −78 °C. Dry tetrahydrofuran (THF) and dichloromethane (CH_2_Cl_2_) were obtained from an Innovative Technology PureSolv system (Inert, Massachusetts, USA). Unless otherwise noted, reactions were performed in oven-dried glassware under an atmosphere of dry argon.

#### Plant Material

The stem bark of *C. fragrans* was collected by L.H.R. in November 2015 in Boina region (Madagascar) and identified by comparison with the authentic sample in the Herbarium of PBZT (Parc Botaniqueet Zoologique de Tsimbazaza, Antananarivo, Madagascar). A voucher specimen of the bark (LivCF2016) was deposited at the College of Pharmacy, The Ohio State University (Columbus, Ohio).

#### Extraction and Isolation

The air-dried stem bark of *C. fragrans* was pulverized and the powder (400 g) was extracted with dichloromethane for 5 days at room temperature. The extract was filtered and concentrated *in vacuo* to yield a yellow-brown oily residue (80.68 g, 20.2% on dry plant material). The residue was divided into fractions using column chromatography over silica gel, eluting with a gradient system of hexanes–EtOAc (from 4:1 to 0:1). Cinnamodial (**1**) was recrystallized using hexanes–EtOAc (1:1) as colorless crystals (5.28 g, 1.32 % on dry plant material).

#### Synthesis

##### Cinnamodiacid (5)

Sodium phosphate monobasic monohydrate (11 mg, 0.08 mmol) in 0.2 mL of water and H_2_O_2_ (66 μL, 0.64 mmol) were added to a solution of **1** (50.0 mg, 0.162 mmol) dissolved in 2 mL of acetonitrile. Sodium chlorite (73.5 mg, 0.8 mmol) in 0.5 mL of water was added dropwise over a period of 15 min. The reaction mixture was stirred for 24 h at room temperature. Solid sodium sulfite, Na_2_SO_3_ (5 mg) was added and the mixture was stirred for 5 min then acidified by an aqueous solution of 10% HCl. Aqueous layer was washed three times with CH_2_Cl_2_. Combined organic layers were washed with brine, dried over MgSO_4_, filtered, and concentrated *in vacuo*. Crystallization in chloroform afforded diacid **5** (43.98 mg, 81%) as colorless crystals; 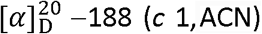; UV (ACN) λ_max_ (log ε 3.69) 204.5 nm; IR (neat, thin film) v_max_ 3435, 3172, 2951, 2929, 2871, 1738, 1395, 1371, 1236, 1073, 1031, 957 and 756 cm^-1^; ^1^H NMR (400 MHz, MeOH) δ 6.86 (1H, d, *J* = 5.1 Hz, H-7), 5.74 (1H, t, *J* = 4.8 Hz, H-6), 2.12 (3H, s, H-17), 2.10 (1H, d, *J* = 4.7 Hz, H-5), 2.09 (1H, td, *J* = 13.7, 3.8 Hz, H-1α), 1.72 (1H, qt, *J* = 13.5, 3.1 Hz, H-2ß), 1.55 (1H, dquint, *J* = 13.7, 3.4 Hz, H-2α), 1.43 (1H, dq, *J* = 12.8, 2.7 Hz, H-3ß), 1.32 (1H, td, *J* = 13.1, 3.1 Hz, H-3α), 1.27 (1H, dm, *J* = 13.8 Hz, H-1ß), 1.29 (3H, s, H-15), 1.22 (3H, s, H-14), 1.04 (3H, s, H-13); ^13^C NMR (100 MHz, MeOD) δ 177.1 (C, C-11), 172.8 (C, C-16), 170.3 (C, C-12), 137.2 (CH, C-7), 136.8 (C, C-8), 79.6 (C, C-9), 68.1 (CH, C-6), 46.3 (CH_2_, C-3), 46.1 (CH, C-5), 42.7 (C, C-10), 35.5 (C, C-4), 34.5 (CH_2_, C-1), 34.2 (CH_3_, C-13), 26.0 (CH_3_, C-14), 22.3 (CH_3_, C-17), 21.2 (CH_3_, C-15), 20.1 (C, C-2); HRESIMS *m/z* 363.14104 [M + Na]^+^ (calcd for C_17_H_24_O_7_Na, 363.14147).

##### Cinnamodiester (6)

To a solution of **5** (50.6 mg, 0.149 mmol) in 1 mL methanol–diethyl ether (1:1), (Trimethylsilyl)diazomethane (80.0 mg, 0.700 mmol, 2.0 M) in hexanes (0.350 mL) was added dropwise at 0 °C. The reaction mixture was stirred at 0 °C for 15 min and concentrated under vacuum to yield diester **6** as an orange amorphous solid: 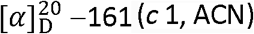; IR (KBr) v_max_ 3501, 2949, 2927, 2868, 1738, 1456, 1371, 1233, 1077, 1033, 754, and 700 cm^-1^; ^1^H NMR (400 MHz, CDCl_3_) δ 6.98 (1H, d, *J* = 5.4 Hz, H-7), 5.72 (1H, t, *J* = 5.4 Hz, H-6), 3.74 (6H, s, (-OCH_3_)_2_) 2.08 (3H, s, H-17), 2.03 (1H, d, *J* = 4.3 Hz, H-5), 1.99 (1H, td, *J* = 13.6, 4.6 Hz, H-1α), 1.61 (1H, qt, *J* = 13.4, 3.2 Hz, H-2ß), 1.52 (1H, dquint, *J* = 13.9, 4.0 Hz, H-2α), 1.38 (1H, dm, *J* = 12.8 Hz, H-3ß), 1.29 (1H, td, *J* = 12.8, 3.0 Hz, H-3α), 1.19 (3H, s, H-15), 1.17 (1H, dm, *J* = 12.8 Hz, H-1ß), 1.14 (3H, s, H-14), 1.00 (3H, s, H-13); ^13^C NMR (100 MHz, CDCl_3_) δ 174.6 (C, C-11), 170.5 (C, C-16), 167.1 (C, C-12), 136.9 (CH, C-7), 133.4 (C, C-8), 78.2 (C, C-9), 66.0 (CH, C-6), 52.8 (OCH_3_ of C-12), 52.5 (OCH_3_ of C-11), 44.4 (CH_2_, C-3), 44.3 (CH, C-5), 41.4 (C, C-10), 34.1 (C, C-4), 33.1 (CH_3_, C-13), 32.9 (CH_2_, C-1), 25.0 (CH_3_, C-14), 21.9 (CH_3_, C-17), 20.2 (CH_3_, C-15), 18.4 (C, C-2); HRESIMS *m/z* 391.17307 [M + Na]^+^ (calcd for C_19_H_28_O_7_Na, 391.17272).

##### Cinnamo-N,11-dihydro-11-pyridazinol (7)

To a solution of **1** (50.0 mg, 0.162 mmol) in 0.5 mL CH_2_Cl_2_–EtOH (1:1) was added to a round-bottomed flask containing a solution of H_2_NNH_2_ • H_2_O (0.015 mL, 0.486 mmol) in 0.16 mL of EtOH at 0 °C via syringe pump. The reaction was warmed to room temperature and was allowed to proceed for 2 h. The reaction mixture was concentrated via rotary evaporation. Toluene (1 mL) was added to the crude material and the resulting mixture was stirred for 10 min at room temperature. The solvent was then removed by rotary evaporation and the process was repeated with CH_2_Cl_2_ (1 mL), yielding **9** as a yellow powder: 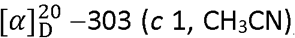; UV (CH_3_CN) λ_max_ (log ε) 286 (3.91), 249 (3.89) nm; IR (KBr) v_max_ 3325, 2949, 2922, 2868, 1735, 1462, 1445, 1370, 1241, 1216, 1081, and 1029 cm^-1^; ^1^H NMR (400 MHz, CD_3_CN) δ 6.89 (1H, s, H-12), 6.06 (1H, s, 9-OH), 5.79 (1H, d, *J* = 5.0 Hz, H-7), 5.67 (1H, t, *J* = 4.8 Hz, H-6), 4.64 (1H, s, H-11), 3.63 (1H, br s, NH), 3.11 (1H, s, 11-OH), 2.13 (1H, d, *J* = 4.6 Hz, H-5), 2.028 (1H, td, *J* = 12.8, 4.1 Hz, H-1α), 2.025 (3H, s, H-17), 1.83 (1H, dq, *J* = 13.6, 2.8 Hz, H-1ß), 1.68 (1H, qt, *J* = 13.7, 3.3 Hz, H-2ß), 1.48 (1H, dquint *J* = 14.1, 3.5 Hz, H-2α), 1.37 (1H, dq, *J* = 12.8, 2.8 Hz, H-3ß), 1.274 (3H, s, H-15), 1.271 (1H, td, *J* = 13.1, 3.2 Hz, H-3α), 1.17 (3H, s, H-14), 0.99 (3H, s, H-13); ^13^C NMR (100 MHz, CD_3_CN) δ 171.7 (C=O, C-16), 140.9 (CH, C-12), 135.3 (C, C-8), 126.5 (CH, C-7), 82.2 (CH, C-11), 73.6 (C, C-9), 68.2 (CH, C-6), 46.5 (CH, C-5), 45.8 (CH_2_, C-3), 41.4 (C, C-10), 35.1 (C, C-4), 34.8 (CH_2_, C-1), 33.8 (CH_3_, C-13), 25.7 (CH_3_, C-14), 22.3 (CH_3_, C-17), 19.7 (C, C-2), 19.1 (CH_3_, C-15); HRESIMS *m/z* 323.19760 [M + H]^+^ (calcd for C_17_H_27_O_4_N_2_, 323.19653).

##### 7β-hydroxy-cinnamopyridazine (18)

^1^H-NMR analysis showed that compound **7** gradually converted to **18** upon incubation in deuterated MeOH at 0 °C overnight (*ca.* 2.5:1 *ratio*, respectively). ^1^H NMR (400 MHz, MeOD) δ 9.24 (1H, s, H-11), 9.04 (1H, s, H-12), 5.82 (1H, br t, *J* = 1.5 Hz, H-6), 4.21 (1H, d, *J* = 1.5 Hz, H-7), 2.43 (1H, dq, *J* = 12.6, 1.9 Hz, H-1ß), 2.06 (3H, s, H-17), 2.01 (1H, dquint *J* = 10.2, 3.7 Hz, H-2β), 1.78 (1H, d, *J* = 0.56 Hz, H-5), 1.74 (1H, m, H-2α), 1.64 (3H, s, H-14), 1.60 (1H, dm, *J* = 13.6 Hz, H-3ß), 1.43 (1H, td, *J* = 12.5, 3.9 Hz, H-1α), 1.38 (1H, td, *J* = 14.0, 3.8 Hz, H-3α), 1.15 (3H, s, H-15), 1.11 (3H, s, H-13); ^13^C NMR (100 MHz, MeOD) δ 172.7 (C=O, C-16), 155.4 (CH, C-12), 151.6 (CH, C-11), 150.7 (C, C-9), 134.3 (C, C-8), 77.7 (CH, C-7), 69.3 (CH, C-6), 49.0 (CH, C-5), 44.4 (CH_2_, C-3), 41.5 (CH_2_, C-1), 38.6 (C, C-10), 35.5 (C, C-4), 34.2 (CH_3_, C-13), 27.0 (CH_3_, C-14), 24.4 (CH_3_, C-15), 22.1 (CH_3_, C-17), 20.7 (C, C-2).

##### **6-*O*-acetyl-12*α*-methyl-pereniporin A (8**) and **6-*O*-acetyl-12*ß*-methyl-pereniporin A (9):**

To a solution of **1** (50 mg, 0.16 mmol) in dry THF (1 mL), methylmagnesium bromide (36.4 mg, 0.486 mmol, 3 eq.) in THF (162 µL) was added dropwise at −78 °C. The reaction mixture was maintained at this temperature, with stirring for 30 m, after which it was allowed to reach r.t. and maintained for another 1.5 h. The reaction mixture was then quenched with NH_4_Cl (1 mL of a saturated aqueous solution). The aqueous solution was extracted with EtOAc (2 mL × 3) and the combined organic layers dried over anhydrous MgSO_4_, filtered, and concentrated. The resulting residue was purified by column chromatography with hexanes–EtOAC (from 3:1 to 2:1) as eluents to afford two diastereomeric compounds (**8** and **9**).

Compound **8** (7.14 mg, 14%) as a colorless oil: 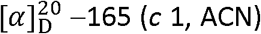; IR (KBr) v_max_ 3427, 2948, 2931, 2869, 1735, 1463, 1372, 1242, 1069, 1024, 983, 960, and 756 cm^-1^; ^1^H NMR (400 MHz, CDCl_3_) δ 5.65 (1H, dd, *J* = 4.0, 1.7 Hz, H-7), 5.61 (1H, td, *J* = 4.4, 1.4 Hz, H-6), 5.27 (1H, s, H-11), 4.41 (1H, qt, *J* = 6.6, 1.6 Hz, H-12), 3.90 (1H, d, *J* = 10.4, 11-OH), 2.05 (3H, s, H-17), 2.03 (1H, d, *J* = 4.0 Hz, H-5), 1.88 (1H, td, *J* = 13.3, 4.4 Hz, H-1α), 1.63 (1H, qt, *J* = 13.5, 3.2 Hz, H-2ß), 1.50 (1H, dquint *J* = 13.8, 3.4 Hz, H-2α), 1.40 (1H, overlapped, H-1ß), 1.39 (3H, d, *J* = 6.6 Hz, 12-CH_3_), 1.38 (1H, overlapped, H-3ß), 1.27 (1H, td, *J* = 13.0, 2.9 Hz, H-3α), 1.16 (3H, s, H-15), 1.15 (3H, s, H-14), 0.99 (3H, s, H-13); ^13^C NMR (100 MHz CDCl_3_) δ 170.7 (C=O, C-16), 146.6 (C, C-8), 120.9 (CH, C-7), 98.1 (CH, C-11), 78.5 (C, C-9), 73.3 (CH, C-12), 67.7 (CH, C-6), 45.9 (CH, C-5), 45.0 (CH_2_, C-3), 38.8 (C, C-10), 34.0 (C, C-4), 33.3 (CH_3_, C-13), 31.9 (CH_2_, C-1), 24.9 (CH_3_, C-14), 23.0 (CH_3_ at C-12), 22.0 (CH_3_, C-17), 19.1 (CH_3_, C-15), 18.3 (C, C-2); HRESIMS *m/z* 347.18280 [M + Na]^+^ (calcd for C_18_H_28_O_5_Na, 347.18290).

Compound **9** (21.26 mg, 40%) as a yellow oil: 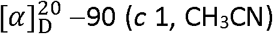; IR (KBr) v_max_ 3434, 2949, 2925, 2868, 1737, 1721, 1462, 1445, 1371, 1242, 1218, 1068, 1022, 986, 958, 913, and 756 cm^-1^; ^1^H NMR (400 MHz, CDCl_3_) δ 5.62 (1H, td, *J* = 4.4, 2.0 Hz, H-6), 5.51 (1H, dd, *J* = 4.0, 2.1 Hz, H-7), 5.39 (1H, s, H-11), 4.77 (1H, qt, *J* = 6.2, 2.1 Hz, H-12), 2.06 (3H, s, H-17), 2.04 (1H, d, *J* = 4.0 Hz, H-5), 1.85 (1H, td, *J* = 13.2, 4.4 Hz, H-1α), 1.64 (1H, qt, *J* = 13.4, 3.1 Hz, H-2ß), 1.50 (1H, dquint *J* = 13.8, 3.4 Hz, H-2α), 1.37 (1H, overlapped, H-3ß), 1.28 (1H, overlapped, H-1ß), 1.27 (1H, overlapped, H-3α), 1.26 (3H, d, *J* = 6.1 Hz, 12-CH_3_), 1.14 (3H, s, H-14), 1.11 (3H, s, H-15), 0.98 (3H, s, H-13); ^13^C NMR (100 MHz, CDCl_3_) δ 170.9 (C=O, C-16), 146.6 (C, C-8), 118.8 (CH, C-7), 96.2 (CH, C-11), 79.3 (C, C-9), 73.3 (CH, C-12), 67.6 (CH, C-6), 45.5 (CH, C-5), 45.0 (CH_2_, C-3), 38.4 (C, C-10), 33.9 (C, C-4), 33.0 (CH_3_, C-13), 32.0 (CH_2_, C-1), 24.8 (CH_3_, C-14), 22.1 (CH_3_, C-17), 18.7 (CH_3_ at C-12), 18.6 (CH_3_, C-15), 18.3 (C, C-2); HRESIMS *m/z* 347.18276 [M + Na]^+^ (calcd for C_18_H_28_O_5_Na, 347.18290).

##### (-)-6*ß*-Acetoxy-9*α*-hydroxydrim-7-ene-12-methyl-12-one-11-al (10)

Compound **9** (*ca*. 20 mg, 0.062 mmol) was dissolved in chloroform and left at room temperature for 22 days, concentrated and purified via column chromatography (hexanes–EtOAc 2:1) to yield **10** (4.49 mg, 22%) as a clear oil: 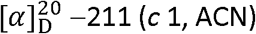; UV (ACN) λ_max_ (log ε 3.37) 218 nm; IR (KBr) v_max_ 3450, 2948, 2930, 2870, 1737, 1673, 1463, 1372, 1233, 1030, 1055, 1030, and 756 cm^-1^; ^1^H NMR (400 MHz, CDCl_3_) δ 9.69 (1H, d, *J* = 0.92 Hz, H-11), 7.00 (1H, d, *J* = 4.9 Hz, H-7), 5.84 (1H, t, *J* = 4.8 Hz, H-6), 4.09 (1H, d, *J* = 1.1, 9-OH), 2.36 (3H, s, 12-CH_3_), 2.14 (3H, s, H-17), 1.99 (1H, d, *J* = 4.5 Hz, H-5), 1.79 (1H, td, *J* = 13.2, 4.5 Hz, H-1α), 1.61 (1H, overlapped, H-2ß), 1.52 (1H, overlapped, H-2α), 1.38 (1H, overlapped, H-3ß), 1.30 (3H, s, H-15), 1.27 (1H, overlapped, H-3α), 1.15 (3H, s, H-14), 1.05 (1H, overlapped, H-1ß), 1.01 (3H, s, H-13); ^13^C NMR (100 MHz, CDCl_3_) δ 200.6 (CH, C-11), 199.8 (C, C-12), 170.6 (C=O, C-16), 141.1 (C, C-8), 140.3 (CH, C-7), 78.3 (C, C-9), 66.7 (CH, C-6), 44.6 (CH, C-5), 44.4 (CH_2_, C-3), 41.7 (C, C-10), 34.2 (C, C-4), 33.0 (CH_3_, C-13), 32.3 (CH_2_, C-1), 25.9 (CH_3_ at C-12), 25.1 (CH_3_, C-14), 21.9 (CH_3_, C-17), 20.2 (CH_3_, C-15), 18.1 (C, C-2); HRESIMS *m/z* 345.16742 [M + Na]^+^ (calcd for C_18_H_26_O_5_Na, 345.16725).

##### Cinnamodial 12-ethylene acetal (11)

To a 10 mL round-bottom flask containing 4 Å molecular sieves, cinnamodial (50 mg, 0.162 mmol) was added and dissolved in anhydrous benzene (6 mL). Ethylene glycol (9.04 µL, 0.162 mmol) and a crystal of *p*-toluenesulfonic acid (1 mg) were added and the solution refluxed for 12 h. The reaction mixture was cooled, ethyl acetate (6 mL) was added and the solution was washed with saturated NaHCO_3_, dried over magnesium sulfate, and evaporated. The residue was purified via column chromatography using a gradient of hexanes–EtOAc (from 4:1 to 2:1) and preparative TLC developed with hexanes–EtOAc (2:1) to afford acetal **11** (3.9 mg, 7%) as fine colorless crystals: 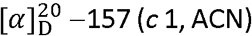; IR (KBr) v_max_ 3485, 2950, 2923, 2871, 1733, 1463, 1390, 1370, 1241, 1213, 1126, 1086, 1062, 1023, 977, 951, 919, and 802 cm^-1^; ^1^H NMR (400 MHz, CDCl_3_) δ 9.87 (1H, d, *J* = 0.72 Hz, H-11), 6.27 (1H, d, *J* = 4.9 Hz, H-7), 5.68 (1H, t, *J* = 4.7 Hz, H-6), 5.17 (1H, s, H-12), 3.90 (2H, m, 12-OCH_2_CH_2_O-), 3.89 (1H, d, *J* = 0.84 Hz, 9-OH), 3.81 (2H, m, 12-OCH_2_CH_2_O-), 2.08 (3H, s, H-17), 2.05 (1H, d, *J* = 4.6 Hz, H-5), 1.87 (1H, td, *J* = 13.3, 4.4 Hz, H-1α), 1.62 (1H, qt, *J* = 13.5, 3.3 Hz, H-2ß), 1.48 (1H, dquint, *J* = 13.8, 3.7 Hz, H-2α), 1.47 (3H, s, H-15), 1.36 (1H, dq, *J* = 13.2, 1.4 Hz, H-3ß), 1.27 (1H, td, *J* = 13.6, 2.9 Hz, H-3α), 1.15 (3H, s, H-14), 0.99 (3H, s, H-13), 0.97 (1H, dm, *J* = 12.8 Hz, H-1ß); ^13^C NMR (100 MHz, CDCl_3_) δ 203.0 (CH, C-11), 170.5 (C, C-16), 136.7 (C, C-8), 131.7 (CH, C-7), 105.5 (CH, C-12), 80.3 (C, C-9), 66.6 (CH, C-6), 65.4 (CH_2_, 12-OCH_2_CH_2_O-), 65.0 (CH_2_, 12-OCH_2_CH_2_O-), 44.9 (CH, C-5), 44.4 (CH_2_, C-3), 41.9 (C, C-10), 34.2 (C, C-4), 32.9 (CH_3_, C-13), 32.9 (CH_2_, C-1), 25.0 (CH_3_, C-14), 22.0 (CH_3_, C-17), 20.4 (CH_3_, C-15), 18.2 (C, C-2); HRESIMS *m/z* 375.17801 [M + Na]^+^ (calcd for C_19_H_28_O_6_Na, 375.17781). Data agree with literature^66^.

##### 12α/ß-methyl-pereniporin A (12)

To a solution of **1** (100 mg, 0.324 mmol) in dry THF (1 mL), methylmagnesium bromide (121.27 mg, 1.621 mmol, 5 eq.) in THF (0.540 mL) was added dropwise at −78 °C. The reaction mixture was maintained at this temperature, with stirring, for 2 h. At this time it was quenched with NH_4_Cl (2 mL of a saturated aqueous solution). The aqueous solution was extracted with EtOAc (4 mL × 3) and the combined organic layers washed with brine (4 mL), dried over anhydrous MgSO_4_, filtered, and concentrated to yield crude **12** (120.68 mg) as a 1:2 diasteromeric mixture of 12α-methyl-pereniporin A and 12ß-methyl-pereniporin A, respectively. IR (KBr) v_max_ 3415, 2972, 2947, 2924, 2869, 1710, 1461, 1383, 1216, 1081, 1065, 1026, 985, 961, and 757 cm^-1^; ^1^H NMR (400 MHz, CDCl_3_) δ (major isomer, C-12*ß*) 5.62 (1H, dd, *J* = 4.0, 2.1 Hz, H-7), 5.38 (1H, s, H-11), 4.79 (1H, qt, *J* = 6.3, 2.3 Hz, H-12), 4.55 (1H, td, *J* = 4.5, 2.3 Hz, H-6), 1.85 (1H, td, *J* = 13.3, 4.5 Hz, H-1α), 1.82 (1H, d, *J* = 5.3 Hz, H-5), 1.66 (1H, qt, *J* = 13.5, 3.3 Hz, H-2ß), 1.51 (1H, dquint *J* = 13.3, 3.9 Hz, H-2α), 1.38 (1H, overlapped, H-3ß), 1.34 (3H, s, H-14), 1.32 (3H, d, *J* = 6.2 Hz, 12-CH_3_), 1.28 (1H, overlapped, H-3α), 1.27 (1H, overlapped, H-1ß), 1.15 (3H, s, H-15), 1.11 (3H, s, H-13); ^13^C NMR (100 MHz, CDCl_3_) δ 144.7 (C, C-8), 123.2 (CH, C-7), 96.4 (CH, C-11), 79.7 (C, C-9), 73.2 (CH, C-12), 66.2 (CH, C-6), 46.7 (CH, C-5), 44.9 (CH_2_, C-3), 38.3 (C, C-10), 34.4 (C, C-4), 33.2 (CH_3_, C-13), 32.5 (CH_2_, C-1), 25.2 (CH_3_, C-14), 19.3 (CH_3_, C-15), 18.9 (CH_3_ at C-12), 18.4 (C, C-2); HRESIMS *m/z* 305.17252 [M + Na]^+^ (calcd for C_16_H_26_O_4_Na, 305.17233).

##### (1′*S/R*)-1′-((8a*S*)-5,5,8a-trimethyl-1,4-dioxo-1,4,4a,5,6,7,8,8a-octahydronaphthalene-2-yl)ethyl formate (13)

To a stirred solution of **12** (100.2 mg, 0.294 mmol, crude, obtained from Grignard addition) in CH_2_Cl_2_ (2 mL) at 0 °C was added PCC (253 mg, 1.175 mmol). The resulting mixture was warmed to room temperature and stirred for 2 h before it was diluted with EtOAc (4 mL) and filtered through a short pad of Celite. The crude mixture was chromatographed on a silica gel column eluting with hexanes–EtOAc (4:1) to obtain **13** (26.77 mg, 27%) as a mixture of diastereomers (*ca.* 1:2 ratio of C-1′*α* and C-1′*ß*, respectively). UV (ACN) λ_max_ (log ε 3.74) 240.5 nm; IR (KBr) v_max_ 2934, 2872, 1729, 1685, 1464, 1380, 1292, 1231, 1166, and 1019 cm^-1^; ^1^H NMR (400 MHz, CDCl_3_) δ (major isomer, C-1′*ß*): 8.02 (1H, d, *J* = 0.49, -OC<u>H</u>O), 6.54 (1H, d, *J* = 1.1 Hz, H-3), 5.72 (1H, qt, *J* = 6.5, 0.92 Hz, H-1′), 2.57 (1H, s, H-4a), 1.85 (1H, dm, *J* = 9.6 Hz, H-8ß), 1.62 (1H, m, overlapped, H-8α), 1.62 (2H, m, H-7ß/α), 1.48 (3H, d, *J* = 6.6 Hz, 1′-CH_3_), 1.43 (1H, dm, *J* = 13.4 Hz, H-6ß), 1.24 (3H, s, 5-ßCH_3_), 1.23 (3H, s, 8a-CH_3_), 1.15 (1H, td, J = 11.2, 4.8 Hz, H-6α), 1.14 (3H, s, 5-αCH_3_); ^13^C NMR (100 MHz, CDCl_3_) δ 203.0 (C, C-4), 199.3 (C, C-1), 160.0 (CH, -O<u>C</u>HO), 148.2 (C, C-2), 135.9 (CH, C-3), 66.4 (CH, C-1′), 61.8 (CH, C-4a), 51.3 (C, C-8a), 42.7 (CH_2_, C-6), 33.6 (C, C-5), 33.5 (CH_2_, C-8), 32.7 (CH_3_, 5-αCH_3_), 21.9 (CH_3_, 5-ßCH_3_), 21.8 (CH_3_, 8a-CH_3_), 20.5 (CH_3_ at C-1′), 17.8 (C, C-7); HRESIMS *m/z* 301.14144 [M + Na]^+^ (calcd for C_17_H_24_O_7_Na, 301.14103).

##### Cysteine methyl ester derivative (16)

Cinnamodial (**1**, 25.5 mg, 0.083 mmol) and *L*-cysteine methyl ester HCL (14.2 mg, 0.83 mmol) were added to an NMR tube, followed by addition of pyridine-*d*_5_ (0.6 mL), vortexed and incubated at 60 °C for 45 min. The mixture was then monitored by NMR at 300.0 K. Compound **16** (98% as observed by ^1^H NMR) was produced after 45 min of incubation. 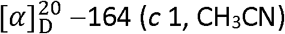; UV (ACN) λ_max_ (log ε 3.72) 250 nm; IR (KBr) v_max_ 3367, 2950, 2928, 2867, 1736, 1677, 1460, 1439, 1371, 1239, 1205, 1167, 1031, and 754 cm^-1^; ^1^H NMR (400 MHz, C_5_D_5_N) δ 5.93 (1H, dd, *J* = 3.9, 1.2 Hz, H-7), 5.90 (1H, br t, *J* = 4.3 Hz, H-6), 5.79 (1H, s, H-12), 5.05 (1H, dd, *J* = 7.2, 1.7 Hz, H-18), 4.77 (1H, s, H-11), 3.54 (1H, s, 19-OCH_3_), 3.50 (1H, dd, *J* = 10.8, 7.1 Hz, H-20a), 3.40 (1H, dd, *J* = 10.8, 2.0 Hz, H-20b), 2.63 (1H, d, *J* = 4.9 Hz, H-5), 2.44 (1H, td, *J* = 13.6, 4.4 Hz, H-1*α*), 1.95 (3H, s, H-17), 1.80 (1H, dm, *J* = 13.6 Hz, H-1*ß*), 1.67 (1H, m, H-2*ß*), 1.48 (1H, dquint, *J* = 13.7, 3.5 Hz, H-2*α*), 1.42 (3H, s, H-15), 1.31 (2H, dm, *J* = 9.2, 2.8 Hz, H-3*α*/*ß*), 1.20 (3H, s, H-14), 0.97 (3H, s, H-13); ^13^C NMR (100 MHz, C_5_D_5_N) δ 172.5 (C, C-19), 170.6 (C, C-16), 144.9 (C, C-8), 122.9 (CH, C-7), 87.6 (CH, C-11), 78.4 (C, C-9), 72.3 (CH, C-12), 68.2 (CH, C-6), 68.1 (CH, C-18), 52.3 (CH_3_, 19-OCH_3_), 45.8 (CH, C-5), 45.4 (CH_2_, C-3), 39.5 (C, C-10), 35.2 (CH_2_, C-20), 34.1 (C, C-4), 33.3 (CH_3_, C-13), 33.0 (CH_2_, C-1), 25.0 (CH_3_, C-14), 21.8 (CH_3_, C-17), 19.7 (CH_3_, C-15), 19.0 (CH_2_, C-2); HRESIMS *m/z* 426.19495 [M + H]^+^ (calcd for C_21_H_32_NO_6_S, 426.19448), 448.17728 [M + Na]^+^ (calcd for C_21_H_31_NO_6_SNa, 426.17643), and base peak is 408.18484 [M – 18 + H]^+^ (calcd for C_21_H_30_NO_5_S).

#### Mosquito cultures and rearing conditions

Eggs of the Liverpool (LVP) strain of *Ae. aegypti* were obtained through the MR4 as part of the BEI Resources Repository, NIAID, NIH (LVP-IB12, MRA-735, deposited by M.Q. Benedict). The eggs of the *Ae*. *aegypti* were reared to adults as described previously^67^.

#### Toxicology assays

Larval and adult toxicities were determined by established protocols^21,68,69^. In brief, for larvae, 10 µl of a derivative (10 mM dissolved in 100% acetone) was added to six wells of a 24-well tissue culture plate containing 1 ml of dH_2_O and five larvae per well, resulting in a final concentration of 100 µM (1% acetone). After 24 h in normal rearing conditions (28°C, 80% relative humidity), the efficacy was assessed by counting the number of larvae per well that did not move after gentle prodding with a micropipette tip or fine insect pin. As a negative and positive control, respectively, the effects of 1% acetone and 100 µM CDIAL were tested in parallel.

For adults, 500 nl of a derivative (3 mM dissolved in 100% acetone) was applied to the thorax of 10 adult females (3-7 days post emergence) to deliver a dose of 1.5 nmol per female. After 24 h in normal rearing conditions, the efficacy was assessed by counting the number of treated mosquitoes that were dead or unable to fly. As a negative and positive control, respectively, the effects of 100% acetone and 1.5 nmol CDIAL were tested in parallel. For both larvae and adult females, the mean efficacies of the derivatives and CDIAL were adjusted for effects of the negative control using Abbott’s correction^56^ and compared statistically using a one-way ANOVA (Bonferroni post-test).

#### Antifeedant assays

A capillary feeding (CAFE) choice assay was used to determine the antifeedant activity of the compounds^21,57,58^. In brief, groups of 5 adult female mosquitoes (3-10 days post-emergence) were placed in *Drosophila* vials (28.5 × 95 mm) covered with cotton plugs. Two 5-µl calibrated glass capillaries were inserted through the cotton into the vial. One capillary was designated the ‘control’ and filled with 5 µl of 10% sucrose containing 1% DMSO (the solvent of the compounds). The other capillary was designated the ‘treatment’ and filled with 5 µl of 10% sucrose containing 1 mM of CDIAL or a derivative (1% DMSO). All vials were placed in normal rearing conditions (28°C, 80% relative humidity) for 18– 20 h after which the volume of sucrose consumed from each capillary was measured. The relative volumes consumed in the treatment vs. control capillaries were used to calculate an antifeedant index^21,57,58^. Each derivative was tested on 5-10 vials of 5 adult females. The mean antifeedant indices of the derivatives and CDIAL was compared statistically using a one-way ANOVA (Bonferroni post-test).

#### Computational Modeling

The modeled structure of the AgTRPA1 monomer (amino acid residues: 537-1191) consists of the last 4 repeats of ankyrin repeat (AR) domain, a transmembrane domain, a linker region between the AR and transmembrane domains, and a C-terminal domain.

Several computational approaches were integrated to build different parts of the AgTRPA1 structure, including *ab initio* modeling^70^, homology modeling, and loop modeling^71^, resulting in a tetrameric structural model of AgTRPA1. This structure was constructed by first forming an AgTRPA1 monomer with GalaxyFill^72^ using the cryo-EM structure of hTRPA1 (PDB ID: 3J9P)^59^ as a template. Subsequently, a tetramer was assembled with GalaxyHomomer^73^ and ZDOCK^74^. Lastly, the tetramer was embedded in a 1-palmitoyl-2-oleoyl-glycero-3-phosphocholine (POPC) bilayer, and the system was minimized/relaxed with short (∼10 ns) molecular dynamics (MD) simulations using CHARMM36 force fields^75^ and GROMACS^76^.

CDIAL was docked to four potential binding pockets near key nucleophilic cysteine and lysine residues (Cys621, Cys641, Cys665 and Lys710 in hTRPA1 and Cys684, Cys704, Lys728 and Lys777 in AgTRPA1) that have been implicated to form covalent bonds with electrophilic agonists^49,50,77^. Among those, the pocket centered around Cys684 in AgTRPA1 was particularly promising. All docking calculations were performed with the Lamarckian genetic algorithm using Autodock 4.2^78^. A 96×68×78 grid box with a grid spacing of 0.375 Å centered around each of the four nucleophilic residues defined the region of the protein that the ligands would explore. 500 docking runs were performed for each AgTRPA1 pocket.

## Acknowledgements

This work was supported in part by the National Institute of Health (NIH) and National Institute of Allergy and Infectious Diseases (NIAID) grant #1R21AI129951-01 to P.M.P and H.L.R. P.K.M gratefully acknowledges support from a Center for Applied Plant Sciences (CAPS) graduate assistantship. P.K.M and H.L.R. are grateful to Dr. W. Tjarks and Dr. J. Fuchs for valuable discussions. P.K.M. and H.L.R. also thank Dr. C. McElroy for assistance during the NMR and MS data collection.

## Author information

### Contributions

P.M.P. and H.L.R. conceived the study. P.M.P., H.L.R. and P.K.M. acquired funding (NIAID and CAPS). P.K.M. and H.L.R. isolated CDIAL, designed, synthesized, and characterized the CDIAL derivatives. M.K. and P.M.P designed, performed, and analyzed the mosquito assays. S.W. and X.C. designed, performed, and analyzed the computational modeling. P.K.M wrote the initial draft of the manuscript. P.K.M., P.M.P., X.C., and H.L.R. contributed to the writing and editing the manuscript. All authors reviewed and approved the final version of the manuscript.

### Corresponding authors

Correspondence to Peter M. Piermarini or Xiaolin Cheng or Harinantenaina L. Rakotondraibe.

